# Metabolic, Biochemical, Mineral and Fatty acid profiles of edible *Brassicaceae* microgreens establish them as promising functional food

**DOI:** 10.1101/2023.05.17.541100

**Authors:** Yogesh Pant, Maneesh Lingwan, Shyam Kumar Masakapalli

## Abstract

Hidden hunger due to micronutrient deficiencies affecting one in three people is a global concern. Identifying functional foods which provide vital health beneficial components in addition to the nutrients is of immense health relevance. Microgreens are edible seedlings enriched with concentrated minerals and phytochemicals whose dietary potential as functional foods needs evaluation. In this study, comprehensive biochemical, mineral, metabolic, and fatty acid profiles of four *Brassicaceae* microgreens - mustard (*Brassica juncea*), pak choi (*Brassica rapa subsp. chinensis*), radish pink (*Raphanus sativus*), and radish white (*Raphanus ruphanistrum*) was investigated. The biochemical and mineral profiling confirmed their promising nutritional and antioxidant nature and as excellent sources of minerals. Mineral profiling using inductively coupled plasma mass spectrometry (ICP-MS) exhibited promising levels of Fe, Mn, Mg, K, and Ca in microgreens. Gas chromatography-mass spectrometry (GC-MS) based metabolite profiling highlighted a range of phytochemicals-sugars, amino acids, organic acids, amines, fatty acids, phenol, and other molecules. Fatty acid profiling established promising levels of Oleic acid (C18:1; Monounsaturated fatty acids-MUFA) and linoleic acids (C18:2; omega-6 Poly unsaturated fatty acid-PUFA), which are health beneficial. It is estimated that fresh microgreens (100 g) can meet about 20 % to 50 % recommended dietary allowance (RDA) of macro- and micro-minerals along with providing useful fatty acids and antioxidants. Overall, the study highlighted *Brassicaceae* microgreens as an excellent nutrient source that can act as functional foods with promising potential to overcome “hidden hunger”.

**Graphical Abstract:** 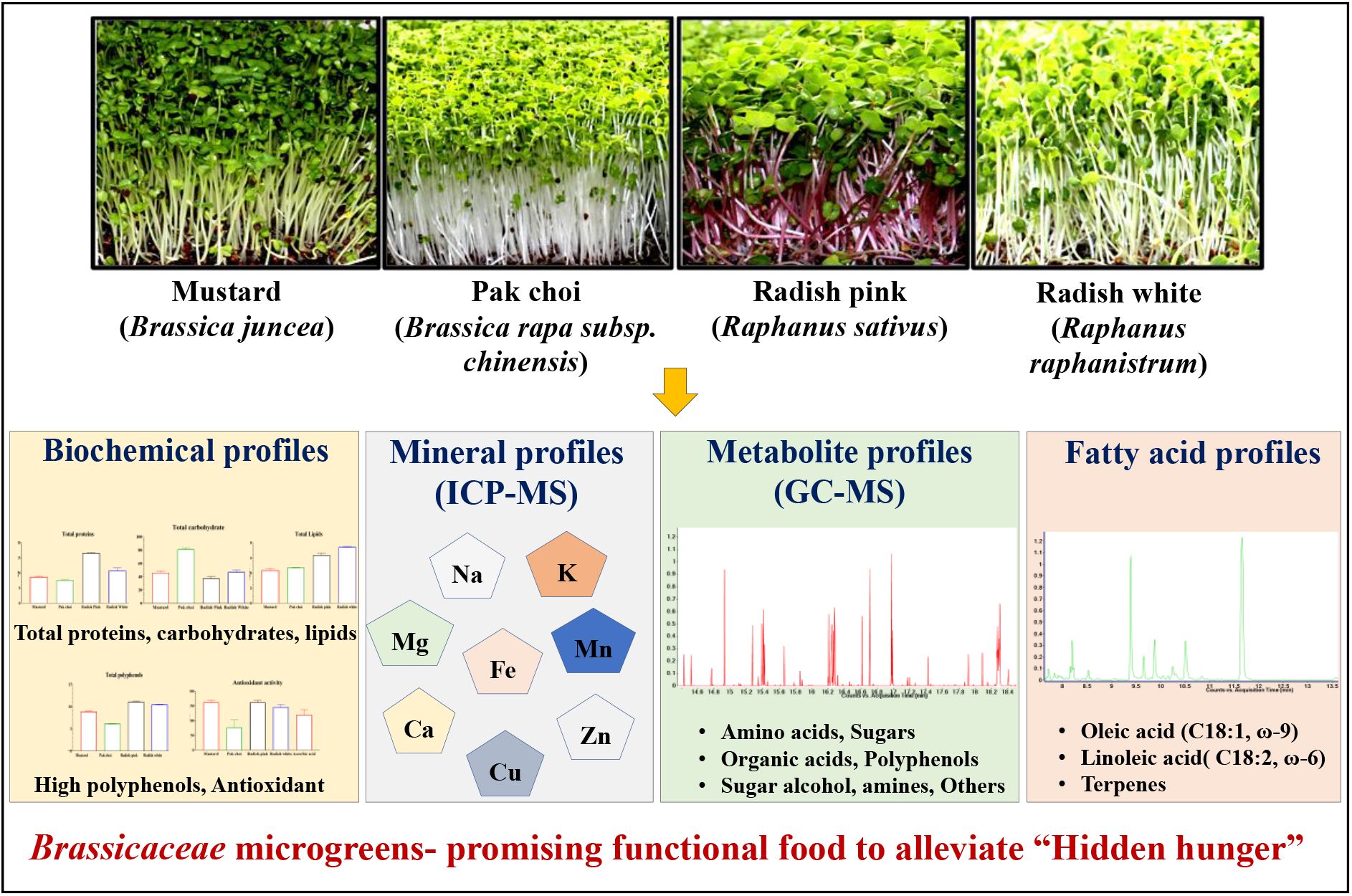

**Highlights:**

- *Brassicaceae* microgreens are rich in molecules with relevance to nutrition and health
- The biochemical analysis supported the antioxidant nature of microgreens
- Comprehensive metabolite profiles of edible microgreens of *Brassica juncea* (Mustard), *Brassica rapa subsp. chinensis* (Pak Choi), *Raphanus sativus* (Radish Pink), and *Raphanus ruphanistrum* (Radish white) using GC-MS are reported
- Ionomics analysis using the *Brassicaceae* microgreens exhibited promising levels of microminerals Fe, Mn, Mg, K, and Ca
- Fatty acid profiles show promising levels of Linoleic acid and Oleic acid, which have health relevance

## 1. Introduction

Despite innovation in global food production, the demand is expected to continuously increase by 35-56 % to feed an approximately 10 billion world population by 2050 (Dijk *et al*., 2021). According to the Food and Agriculture Organization (FAO), 41.9 percent of the world’s population could not afford a healthy diet in 2019 (FAO, 2019). This includes the deficiency in multiple micronutrients while consuming an energy-efficient diet or a minimum number of calories, termed “hidden hunger.” Identifying and adopting micronutrient-rich diets to existing foods will immensely benefit global health. Young edible seedlings, now termed microgreens, are being promoted as nutritionally promising candidates to address the problems of hidden hunger and malnutrition (Bhaswant *et al*., 2023).

Microgreens are young seedlings grown in soil or *in-vitro*-controlled environmental conditions from the seeds of vegetables and herbs, having two fully developed cotyledon leaves. Their harvesting stage depends upon the growing conditions and species type, varying from 7-21 days after germination (Xiao, 2012). The major families contributing to microgreens include *Brassicaceae, Asteraceae, Apiaceae, Amaryllidaceae, Amaranthaceae, Cucurbitaceae, Fabaceae, Poaceae*, and *Lamiaceae*. They are known for both flavoring and nutrition, and numerous studies have established their dietary importance in terms of mineral contents, vitamins, antioxidants, etc. (Lester *et al*., 2010). Studies on twenty-five different varieties of microgreens, such as arugula, celery, cilantro, radish, and amaranth, showed higher concentrations (4 to 40 times) of nutrients, antioxidants, and vitamins than mature plants (Xiao *et al*. 2012).

The *Brassicaceae* family is reported to be rich in antioxidants, phenolics, vitamins, minerals, and other phytochemicals like glucosinolates with anti-inflammatory and anticarcinogenic activity (Bell *et al*., 2017). It is shown that microgreens could fulfill children’s dietary requirements for minerals such as Ca, Mg, Fe, Mn, Zn, Se, and Mo (Pinto *et al*., 2015). Furthermore, low potassium-containing microgreens were recommended for patients with reduced kidney function (Renna *et al*., 2018). Studies in mice suggest that when microgreens are supplemented with a high-fat diet, they can modulate weight gain and cholesterol metabolism and may protect against cardiovascular diseases by preventing hypercholesterolemia (Huang *et al*., 2016). These statements nominate microgreens to be considered a “Superfood.” However, there is still a need for a comprehensive study to evaluate the antioxidant potential, phytochemical, and mineral (ionomics) profiling of *Brassicaceae* microgreens. In addition, it will be interesting to investigate fatty acids in *Brassicaceae* microgreens; being a rich source of oils, the unsaturated fatty acids available in the microgreens will be immensely relevant to health benefits.

This study is focused on biochemical analysis and comprehensive profiling of metabolites, minerals, and fatty acids of four *Brassicaceae* microgreens - mustard (*Brassica juncea*), pak choi (*Brassica rapa subsp. chinensis*), radish pink (*Raphanus sativus*), and radish white (*Raphanus ruphanistrum*). The optimal *in-vitro* cultivation of microgreens under controlled conditions was achieved for the study. The biochemical content of total proteins, carbohydrates, lipids, phenols, and antioxidants is evaluated. The mineral concentrations, metabolic and lipid profiles are investigated using sensitive analytical platforms of ICP-MS, GC-MS, and GC-FAME, respectively. Finally, we compared these profiles with nutritional significance, establishing the *Brassicaceae* microgreens as a promising functional food.

## 2. Material and methods

All the chemicals and reagents were purchased from Sigma Aldrich, unless mentioned in the methods.

### 2.1 Plant Materials and growth conditions for Microgreens

Four commonly consumed microgreens from the family *Brassicaceae* were selected in this study. The microgreen seeds of *Brassica juncea* (Mustard), *Brassica rapa subsp. chinensis* (Pak Choi), *Raphanus sativus* (Radish Pink), and *Raphanus ruphanistrum* (Radish white) were purchased from the local company “AllThatGrows” (https://www.allthatgrows.in/). Further substrate, seed density, and germination parameters were investigated for the efficient growth of these microgreens. In general, the seeds were sown in different combinations of coco-peat and vermiculite in the ratios 1:0, 3:1, 1:1, and 1:3 in a square petri plate (144 cm^2^, HiMedia-PW050) with adequate moisture content. These were kept in dark for three days at 22 ± 2 °C temperature, 60 ± 5 % relative humidity (RH), and the germination percentage was calculated. Finally, the emerged seedlings were transferred to light with a 16/8 hour light-dark cycle, after which the shoots of microgreens (6-9 cm height) were harvested at the fully grown, two-leaf stage (Day 7). The samples were quenched using Liquid N_2_ and stored at -80 °C for further analysis.

### 2.2 Nutritional Analysis in Microgreens

#### 2.2.1 Estimation of total proteins

Total proteins were estimated according to the standard DC protein assay (Bio-Rad, catalog number 500-0116). For protein extraction, 20 mg of fresh sample was weighed and crushed in liquid nitrogen. Further, 200 μl of extraction buffer (40 mM Tris-HCl, 250 mM Sucrose, 10 mM EDTA) was added to each vial (Wu *et al*., 2014). The content was then vortexed for 20 sec and incubated in ice for 5 min. This step was repeated thrice for 15 min, followed by centrifugation at 15,000 rpm for 20 min at 4°C. Next, the supernatant was collected and used for the assay. Next, 5 μl of this aliquot was taken, and 25 μl Reagent A’ (an alkaline copper tartrate solution) was added. Further, 200 μl of Reagent B (a dilute Folin Reagent) was added, and the content was mixed slowly. After 15 min of incubation in the dark, the absorbance of the sample was taken at 750 nm in a microplate reader. Total proteins were estimated against BSA (Bovine serum albumin) standard curve.

#### 2.2.2 Estimation of total lipid

Lipid extraction and quantification were done using the gravimetric approach (Bligh and Dyer 1959). Freshly harvested microgreens (1 gm) were crushed with liquid nitrogen, and 3 ml of chloroform: methanol (1:2) was added. The samples were homogenized and centrifuged at 3,000 rpm for 5 min. The supernatant was collected separately, and 3 ml of chloroform: methanol (1:2) with 0.8 ml of 1 % KCl was added to the pellet. The above centrifugation step was repeated, and both supernatants were pooled together. Further, 2 ml of chloroform and 1.2 ml of 1 % KCl was added to it and vortexed, followed by centrifugation at 3,000 rpm for 5 min. Finally, the bottom layer was collected from the pooled lipid extract sample into a previously weighed centrifuge tube and subjected to solvent evaporation. The final weight of the tubes was recorded, and total lipids were calculated as an increase in weight.

#### 2.2.3 Estimation of total carbohydrates

20 mg of fresh microgreens sample were ground in a pestle and mortar using liquid nitrogen. Carbohydrate extraction was performed in 80 % ethanol (1 ml) (Bauer *et al*., 2022). The samples were vortexed and centrifuged at 15,000 rpm for 10 min. The supernatant was collected, and 4 ml of anthrone reagent (2 mg/ml in H_2_SO_4_) was added. The mixture was again vortexed and placed in a heat block at 100 °C for 10 min. The tubes were allowed to cool at room temperature, and absorbance was recorded at 630 nm. Total carbohydrates were calculated using glucose as standard.

#### 2.2.4 Estimation of total phenolic compounds

Fresh microgreens (100 mg) were crushed in liquid nitrogen and mixed with 70 % acetone (2 ml) for phenols extraction. The samples were vortexed and centrifuged at 10,000 rpm for 10 min at 4 °C. The procedure was repeated with 1 ml of 70 % acetone, and the supernatants were pooled. Next, 100 μl of this sample was mixed with 500 μl Folin-Ciocalteau reagent (10% v/v in MiliQ water) and incubated for 5 min (Ainsworth and Gillespie 2007). The reaction was initiated by adding 400 μl sodium carbonate (5 % w/v in water) and setting it for 20 min in the dark at room temperature. Total soluble phenols were calibrated using Gallic acid as a standard (concentration ranging from 20-100 μg/ml) after absorbance was measured at 765 nm with a spectrophotometer against water as blank.

#### 2.2.5 Determination of DPPH radical-scavenging activity

500 mg of fresh-weight of samples were taken and crushed in liquid nitrogen. 10 ml of 80 % ethanol was added, and the content was centrifuged at 10,000 rpm for 15 min. The supernatant was collected in a separate vial. 900 μl of DPPH solution (0.1 mM DPPH in 80 % ethanol) was mixed with 100 μl of different concentrations of sample extract (five concentrations-0.04 % to 0.2 %). The reaction mixture was vortexed and kept at room temperature in the dark for 30 min. The decrease in absorbance was recorded at 515 nm (Lingwan *et al*., 2021). Inhibition percent (I %) of the free radical DPPH• in microgreens samples was expressed as-I % = (A_control_ - A_sample_) x 100/A_control_; where: A_control_ is the absorbance of blank DPPH solution; A_sample_ is the absorbance of samples. Finally, IC50 (50 % inhibitory concentration) was calculated, and values were compared to a positive ascorbic acid standard. The less the IC50 value to ascorbic acid (0.12 μg/ml), the more it is antioxidant.

### 2.3 Elemental analysis through ICP-MS

Powdered microgreens (100 mg dry weight) were subjected to acid digestion with 3.2 ml nitric acid and 800 μl of hydrogen peroxide (Zou *et al*., 2021). The samples were filtered using a 0.2-micron filter and diluted five times with MilliQ water. Finally, the samples were subjected to ICP-MS along with standards for quantification of the following elements-Na, Mg, K, Ca, Mn, Fe, Cu, and Zn. The quantities were expressed in milligrams per gram dry weight (mg/g DW) for macro minerals and microgram per gram dry weight for micro minerals (μg/g DW). Further, the recommended dietary allowance (RDA) % was calculated as the percent contribution of minerals present in microgreens to the recommended levels by *FSSAI*.

### 2.4 Gas chromatography-mass spectrometry (GC-MS) based profiling of metabolites

Metabolite extraction was performed in lyophilized microgreens samples (25 mg) with 1.2 ml of 80 % methanol (Lisec *et al*. 2006). 20 μl of Ribitol (0.01 % w/v) was added to each sample as an internal standard for relative quantification. These were incubated in a thermomixer at 70 °C and 900 rpm for 5 min and centrifuged at 13,000 rpm for 15 min at 25 °C. Next, 50 μl of supernatant was dried in a speed vac. for further derivatization step. 35 μl of pyridine containing methoxyl amine hydrochloride (20 mg/ml) was added to each dried sample and incubated at 37 °C, 900 rpm for 2 hours. Later, 49 μl of MSTFA (N-methyl-N- (trimethylsilyl)- trifluoroacetamide) was added to the tubes and incubated for 30 min. The sample was centrifuged at 13,000 g for 10 min, and the supernatant was transferred to new inserts (0.2 ml volume) for GC-MS data acquisition (Masakapalli *et al*., 2014; Lingwan and Masakapalli, 2022).

GC-MS-based analysis was performed using Agilent Technology GC, model no. 7890B with a run time of 60 min in splitless mode using helium as carrier gas at a flow rate of 0.6 ml/min. The program was set initially at 50 °C temperature for 1 min, increasing to 200 °C for 4 min at the rate of 10 °C and finally to 300 °C at 5 °C/min for 10 min (Shree *et al*. 2019). The mass spectra were processed through Metalign software for baseline correction. Further, analyzing retention time and fragmentation patterns, metabolites corresponding to the peaks were identified through MassHunter Qualitative Navigator software using NIST version 2.3, 2017, and Fiehn Metabolomics 2013 libraries with an identity score of ≥70 %.

### 2.5 Lipid extraction and GC-FAME-based profiling

For lipid extraction and transesterification, fresh microgreens samples (50 mg) were crushed in liquid nitrogen and saponified with 1 ml of saturated methanolic KOH at 100 °C for 30 min. After 2 min incubation at room temperature, 2 ml of 5 % HCl prepared in methanol was added to the extract and subjected to 80 °C for 10 min. Additionally, 1.25 ml solution of 1:1 n-hexane and methyl tertiary-butyl ether was added, and the mixture was gently mixed. The tubes were positioned upright for phase separation, and the top layer was collected and washed with 3 ml of 1.2% KOH solution. Finally, a saturated NaCl solution was added to completely separate the n-hexane phase containing Fatty acid methyl esters (FAMEs) (Woo *et al*., 2012). 1 μl of these extracts were directly injected into the GC-MS equipped with HP-5 (30 m, 0.32 mm i.d., 0.25 μm) column. The method parameters include a run time of 40 min in splitless mode using helium as carrier gas at a flow rate of 1 ml/min. The program was set initially at 50 °C temperature for 1 min, increasing to 200 °C at 25 °C/min for 5 min and finally to 230 °C at 3 °C/min for 18 min.

### 2.6 Statistical and multivariate data analysis

The identified compounds and their respective abundances were subjected to multivariate statistical analysis using MetaboAnalyst 5.0 online tool. Data pre-processing was performed by normalization by the median, log transformation, and Pareto scaling. Finally, PCA (Principal component analysis), PLS-DA (Partial least square discriminant analysis) plots, and Heat maps were generated (Chong *et al*., 2019). The area of the internal standard, Ribitol, is used to obtain the relative proportions of the peaks, leading to the calculation of fold changes of metabolite/peak levels among the treatments. Fundamental statistical methods were used to determine the significance or non-significance of data using GraphPad Prism8 software.

## 3. Results and discussion

### 3.1 Optimal growth of *Brassicaceae* microgreens

Substrate, Seed density, and germination percentage were optimized for the efficient growth of microgreens. Different ratios of Coco-peat: vermiculite (1:0, 3:1, 1:1, and 1:3) were tested as substrates for microgreens. Based on the maximum number of seeds germinated, the 1:1 ratio of coco-peat and vermiculite mixture was observed to be the best composition. The seeds of selected microgreen seeds were categorized under three seed densities (low, medium, and high) based on their relative sizes. Seed density of 2.5 g and 3 g for pakchoi and mustard, and 3.5 g for radish pink and radish white per 144 cm^2^ (grown in square petri plate) was found to be optimum. The germination percentage of four selected microgreens was between 86.5 ± 9 % and 98 ± 4%. There were no significant differences among the microgreens since all germinated efficiently on the cocopeat-vermiculite (1:1) substrate (Figure 1A). Microgreens can grow on multiple substrates, jute mats, potting mixes, and hydroponically. Therefore, optimal substrate combinations are vital for producing safe and high-yield microgreens, and a coco-peat and vermiculite mixture could be considered.

**Figure 1.**
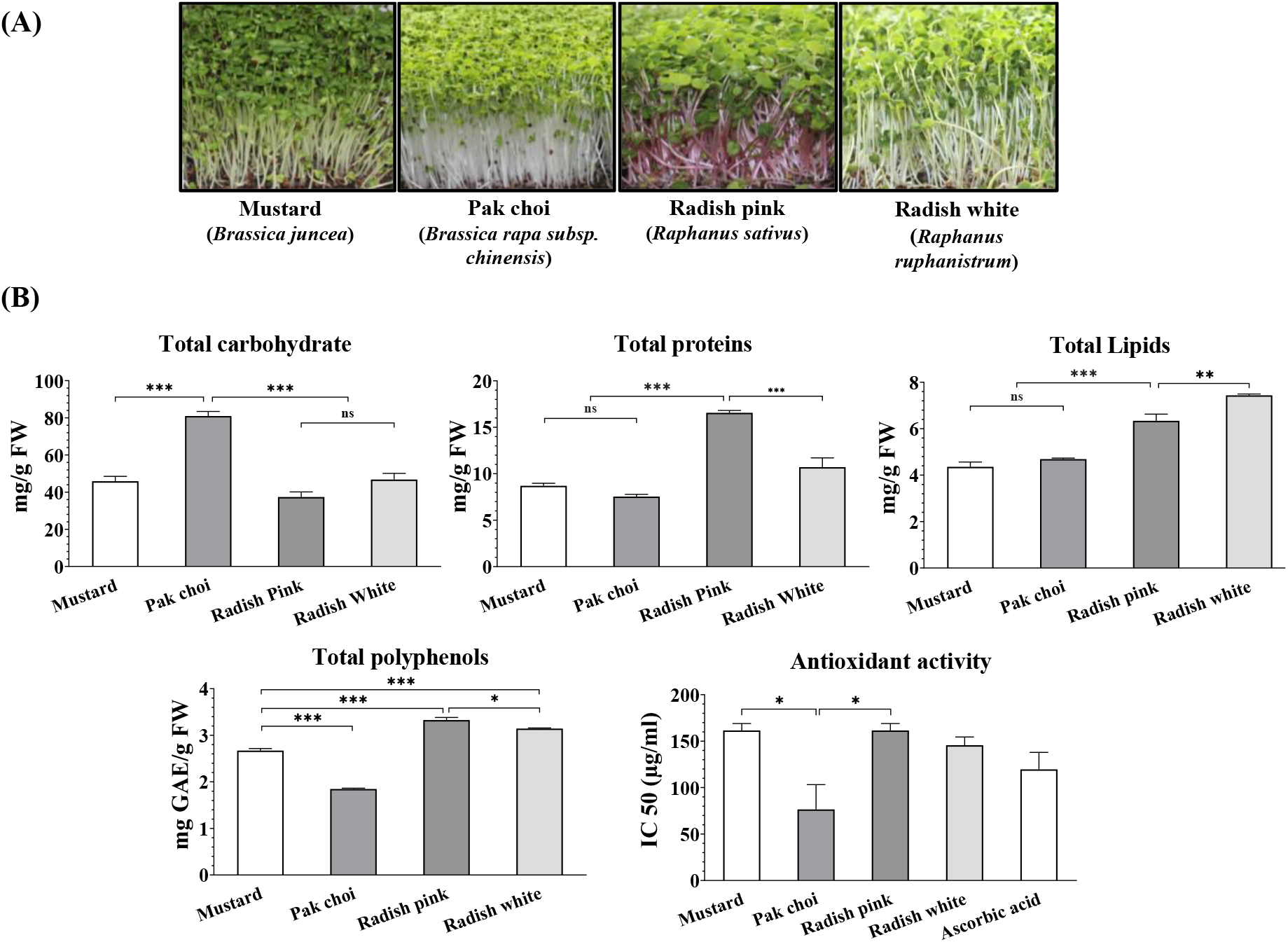
(A) 7 days old microgreens of Mustard, Pak choi, Radish Pink, and Radish white grown on 1:1 cocopeat-vermiculite substrate. (B) Comparison of total carbohydrates, proteins, lipids, polyphenols, and antioxidant activity in *Brassicaceae* microgreens expressed in mg g^-1^ fresh weight (FW). Error bars represent the standard deviation for n=3

### 3.2 Biochemical analysis show higher nutritional and antioxidant nature of *Brassicaceae* microgreens

The study of biochemical analysis of *Brassicaceae* microgreens shed light on their promising nutritional and antioxidative potential. The total carbohydrates ranged between 32 and 86 mg/g of fresh weight (FW) in the microgreens (Figure 1B). No apparent differences were found among mustard, radish pink, and radish white microgreens and are similar to the levels present in mature mustard leaves (U.S. Department of Agriculture, 2019). Pak choi microgreens showed 2 to 4-fold higher carbohydrate levels than the others (U.S. Department of Agriculture, 2019; Goyeneche *et al*., 2015). The total proteins were observed to be ranging from 7 mg/g FW to 17 mg/g FW in microgreens which were less than the total proteins reported in mature leaves of mustard, pak choi, and radish, i.e., 15 to 40 mg/g FW (Goyeneche *et al*. 2015; U.S. Department of Agriculture, 2019). The total lipids ranged between 3.9 mg/g FW and 7.5 mg/g FW. No significant differences were observed between mustard and pak choi microgreens. Also, both radish pink and radish white varieties had similar lipid content. However, the lipids were less in mustard and pak choi than in radish greens (Figure 1B). Furthermore, these levels were equivalent to the mature *Brassicaceae* leaves (Goyeneche *et al*., 2015; U.S. Department of Agriculture, 2019).

The total polyphenols and DPPH scavenging activity of *Brassicaceae* microgreens established their promising antioxidant potential (Figure 1B). The total polyphenol content (TPC) in microgreens ranged from 1.85 mg to 3.33 mg GAE/g FW which were higher compared to the whole mustard plant (3.5 mg/g dry weight) and mature leaves (17.7 mg/g dry weight) reported in the literature (Sun *et al*., 2018). We observed promising antioxidant potential of *Brassicaceae* microgreens as evidenced by DPPH radicle scavenging abilities with IC50 of 76.5 μg/ml to 161.5 μg/ml compared to the positive control ascorbic acid (IC50: 119.6 g/ml). Comparatively, similar values were noted in other microgreens previously reported (Ghoora *et al*., 2020). An antioxidant-rich diet reduces the risk of cardiovascular diseases, hypertension, and diabetes (Alissa and Ferns, 2017). In addition, polyphenols and other antioxidants function as scavengers and reduce oxidative stress, which helps manage chronic non-communicable diseases (Urquiaga and Leighton, 2000). Overall, TPC was higher in microgreens than in mature plants, which can directly correlate to healthy human nutrition.

### 3.3. *Brassicaceae* microgreens are an excellent source of minerals

Minerals are necessary for any organism to function correctly and are vital to the human diet. They regulate physiological and biochemical activities, metabolism, and homeostatic balance (Kyriacou *et al*., 2021). Here we evaluated the *Brassicaceae* microgreens as a potential source of minerals (macro and micro) measured through ICP-MS analysis (Table 1). Macro minerals Na and K were found in concentrations ranging from 17 to 27 mg/g DW and 36 to 61 mg/g DW respectively. In comparison, Ca and Mg were present at 9 to 17 mg/g DW and 6 to 9 mg/g DW. Among the microminerals, Fe has the highest concentration (277 to 1092 μg/g DW), while Cu has the lowest concentration (7.5 to 10.4 μg/g DW). Additionally, Mn and Zn are present in the range between 53.5 and 206.5 μg/g DW of microgreens. The data was compared to the *Food Safety and Standards Authority of India* (FSSAI) 2020 recommendations for recommended dietary allowance (RDA). It is known that fruit and vegetable intake supply approximately 11% Na, 24% Mg, 35% K, 7% Ca, 21% Mn, 16% Fe, 30% Cu, and 11% Zn of recommended RDA to the human body (Levander, 1990). From the elemental profile, it is clear that all the selected *Brassicaceae* microgreens are excellent sources of macro and micro minerals. Consuming mustard microgreens (about 10 grams DW) can fulfill 50 % RDA of iron (Table 1). Overall, these *Brassicaceae* microgreens can provide 10-20 % of the RDA for macronutrients and 4-50 % of the requirement for micronutrients, depending on the mineral type and species. The analysis confirmed that the levels of Fe, Mn, Mg, K, Ca, and Na in *Brassicaceae* microgreens were higher when compared with the mineral content of mature wild edible parts of mustard, pak choi, and radish (Filho *et al*., 2018, Kyriacou *et al*., 2021, Mezeyova *et al*., 2022). High levels of minerals have also been reported in other *Brassicaceae* microgreens, such as arugula, broccoli, and red cabbage, compared with their mature leaves (Johnson *et al*., 2021; Supplementary table S2). In addition, calcium and magnesium, critical elements in the human diet, were higher in all the *Brassicaceae* microgreens studied (Armesto *et al*., 2019).

**Table 1.** Different macro and micro minerals concentrations in *Brassicaceae* microgreens measured using ICP-MS. The estimated RDA % of minerals met via dietary source on consuming 10 g DW of microgreens daily (equivalent to 100 g FW) is tabulated and described as the percent contribution of minerals present in *Brassicaceae* microgreens to recommended dietary allowance (RDA) values approved by *FSSAI* 2020.

| Macro-minerals (mg/g DW) |  |  |  |  |  |  |  |  |  |
| --- | --- | --- | --- | --- | --- | --- | --- | --- | --- |
|  | RDA<br>(mg per day),<br>by FSSAI | Mustard | RDA %* | Pak choi | RDA %* | Radish<br>pink | RDA %* | Radish<br>white | RDA %* |
| Na | 2000 | 20.7 ± 1.44 | 10 | 26.3 ± 1.33 | 13 | 20.9 ± 1.99 | 10 | 18.2 ± 1.03 | 9 |
| Mg | 385 | 7.3 ± 0.52 | 19 | 8.6 ± 0.38 | 22 | 6.6 ± 0.70 | 17 | 6.9 ± 0.43 | 18 |
| K | 3500 | 56.3 ± 4.0 | 16 | 58.3 ± 3.1 | 17 | 52.9 ± 5.3 | 15 | 38.7 ± 2.5 | 11 |
| Ca | 1000 | 14.4 ± 1.5 | 14 | 16.9 ± 0.64 | 17 | 10.4 ± 0.98 | 10 | 9.4 ± 0.56 | 9 |
| Micro-minerals (µg/g DW) |  |  |  |  |  |  |  |  |  |
|  | RDA<br>(µg per day) |  |  |  |  |  |  |  |  |
| Mn | 4000 | 115.7 ± 8.8 | 29 | 88.5 ± 4.2 | 22 | 51.2 ± 5.1 | 13 | 50.4 ± 3.1 | 13 |
| Fe | 19000 | 1021 ± 71.9 | 51 | 266.4 ± 11.5 | 13 | 624.6 ± 66.1 | 31 | 293.6 ± 18.8 | 15 |
| Cu | 2000 | 7.4 ± 0.9 | 4 | 9.6 ± 0.8 | 5 | 8.0 ± 0.6 | 4 | 6.7 ± 0.8 | 3 |
| Zn | 17000 | 135.9 ± 10.3 | 8 | 196.2 ± 10.3 | 12 | 136.3 ± 13.9 | 8 | 118.9 ± 6.9 | 7 |

\* Percent RDA (%) met after consuming 10 g DW of *Brassicaceae* microgreens [compared with RDA 2020; FSSAI].

Many of these macro and micro minerals are commonly deficient in the population of both developed and developing countries, the symptoms of which are not always immediately visible. Hence, including *Brassicaceae* microgreens in the diet which are observed to be dense in minerals, can assist in meeting daily needs and overcoming “hidden hunger.”

### 3.4. GC-MS-based metabolite profiling shows microgreens are a rich source of bioactive compounds

The aqueous methanol soluble metabolite profiles of *Brassicaceae* microgreens using GC-MS captured sugars, amino acids, fatty acids, organic acids, and bioactive compounds, including various polyphenols, sugar alcohols, and amines. (Figure 2, Supplementary Table S1). Among the sugars and sugar alcohols, Fructose, glucose, meso-erythritol, and threitol were predominant (Supplementary Figure S1). Other sugars identified are mannose, arabinose, glycerol, erythritol, glucopyranoside, and myoinositol. The amino acids; alanine, valine, leucine, isoleucine, glycine, serine, threonine, aspartic acid, glutamic acid, phenylalanine, asparagine, glutamine, lysine, and tyrosine were identified. These include essential amino acids, which can be of nutritional relevance. In addition, the branched-chain amino acids valine, leucine, and isoleucine, generally recommended in protein supplements for athletes, could be interesting. Myristic acid, palmitic acid, linolenic acid, and stearic acid are among the identified fatty acids. Organic acids found in microgreens, such as lactic acid, glycolic acid, citric acid, malic acid, amino-butanoic acid, glyceric acid, and butenedioic acid, may aid in human digestion (Nguyen and Kim, 2020). The metabolite profiles also revealed the presence of polyphenols and amines of nutritional relevance in all the *Brassicaceae* microgreens. These include sinapinic acid, oxoproline, and hydroxylamine. In addition, amino furanone and an unknown metabolite at a retention 22.52 min were also reported. Overall, the metabolite profiles of microgreens showed several small molecules belonging to sugars, amino acids, fatty acids, organic acids, polyphenols, sugar alcohols, and amines that have nutritional significance.

**Figure 2.**
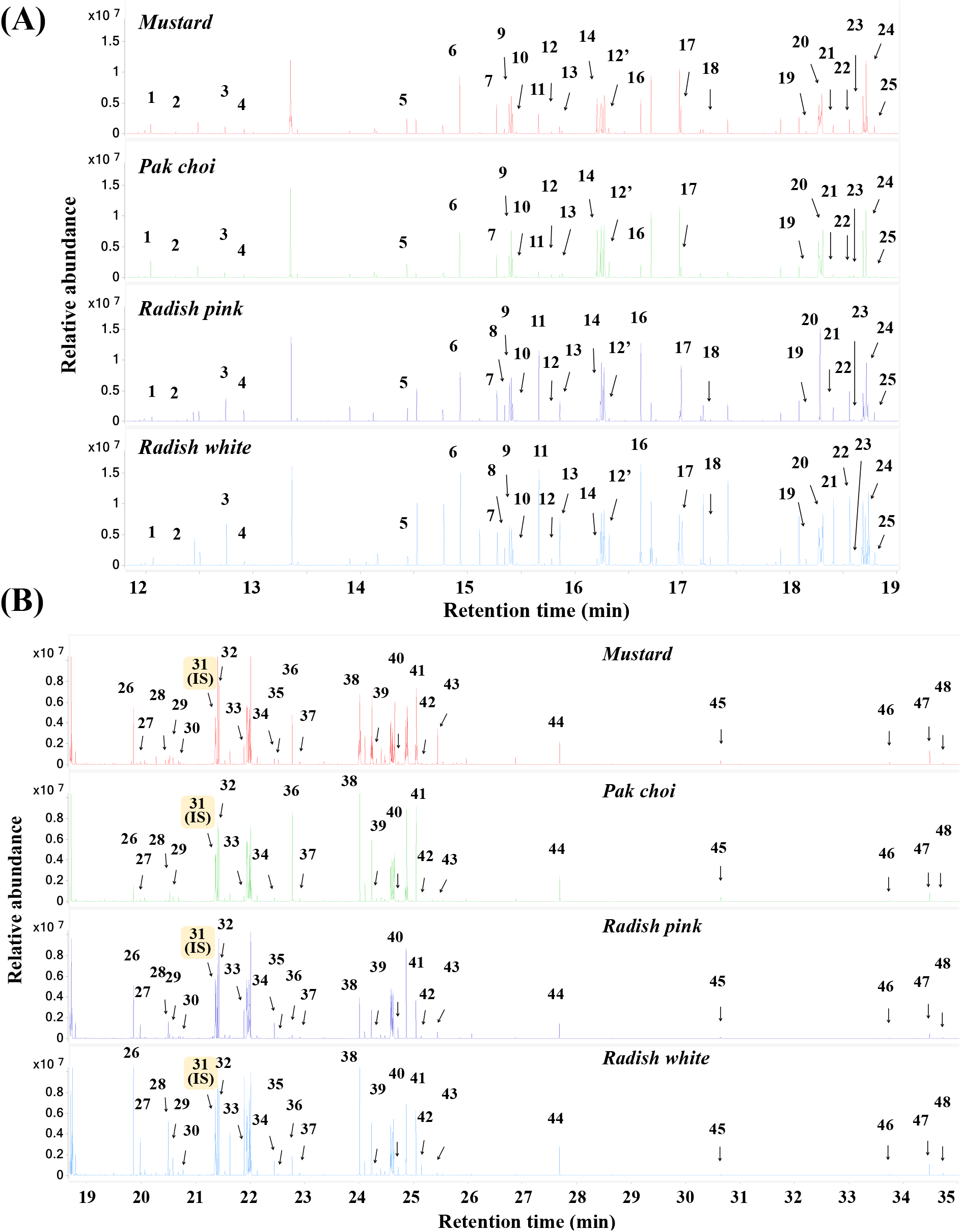
Overlapping total ion chromatograms (TICs) of seven days old *Brassicaceae* microgreens from retention time. (A) 12 min to 19 min and (B) 20 to 35 min, covering all the identified metabolites range. Ribitol was used as an internal standard (IS). The metabolites identified correspond to the number as follows-1. Lactic Acid; 2. Glycolic acid; 3. Alanine; 4. Hydroxylamine; 5. Valine; 6. Benzoic Acid; 7. Ethanolamine; 8. Leucine; 9. Glycerol; 10. 4-Dimethylamino-2(5H)-furanone; 11. Isoleucine; 12. 2-Butenedioic acid; 13. Glycine; 14. Glyceric acid; 15. Nonanoic acid; 16. Serine; 17. Threonine; 18. Aspartic acid; 19. Timonacic; 20. Malic acid; 21. 3-Amino-2-piperidone; 22. Methyl L-alaninate; 23. Erythritol; 24. Oxoproline; 25. 4-Aminobutanoic acid; 26. Glutamic acid; 27. Phenylalanine; 28. Asparagine; 29. Arabinose; 30. Ribono-1,4-lactone; 31. Ribitol; 32. meso-Erythritol; 33. Glutamine; 34. Threitol; 35. Metabolite@22.52; 36. Citric acid; 37. Myristic acid; 38. Fructose; 39. Mannose; 40. Lysine; 41. Glucose; 42. Tyrosine; 43. Glucopyranoside; 44. Palmitic Acid; 45. Myo-Inositol; 46. α-Linolenic acid; 47. Stearic acid; 48. Sinapinic acid. The spectra in red, green, dark blue, and sky blue correspond to mustard, pak choi, radish pink, and radish white, respectively.

### 3.5. Multivariate statistical analysis showed an overall variation among the microgreen species

The variations in the metabolite profiles among the *Brassicaceae* microgreens were further captured via Multivariate statistical analysis. Principal component analysis (PCA) showed distinct clusters of microgreens represented by the first two principal components (PCs), where PC1 and PC2 explained 65.4% and 12.9% of the variance in metabolite profiles, respectively (Figure 3A). Additionally, the relative abundances of identified metabolites among the microgreens were presented in bar graphs (Supplementary figure S2). This confirms that the composition of soluble metabolites among the microgreens is distinct, which could contribute to different attributes such as taste, color, smell, texture, etc., along with other parameters.

**Figure 3.**
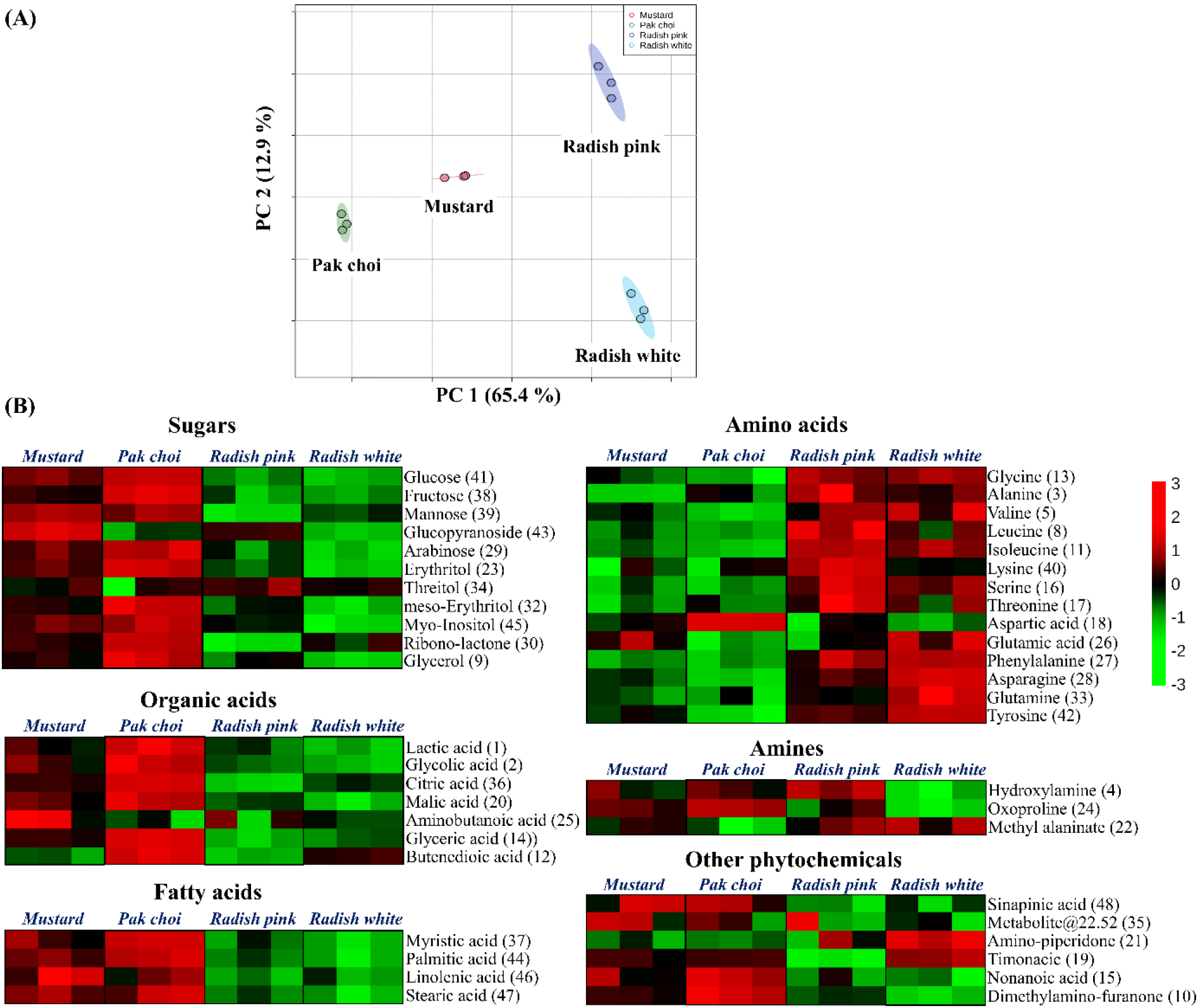
Microgreens from the *Brassicaceae* family have distinct metabolic profiles. A) Principal Component Analysis (PCA) plots depict the variation among the *Brassicaceae* microgreens. B) The heat map shows the metabolic response to microgreens, characterized as sugars, amino acids, organic acids, amines, fatty acids, and other phytochemicals. For each of the observed metabolites, the average relative abundance (n=4) is shown as a set of color-coded groups, from red (+3, high) to green (−3, low). The numbers against each metabolite correspond to the designated peaks observed in the GC-MS spectra depicted in Figure 2.

The heat map exhibited the response of microgreens’ metabolites, classified as sugars, amino acids, organic acids, amines, fatty acids, and other phytochemicals. Results indicate that the metabolite profiles of mustard and pak choi are comparable with higher levels of identified sugars, organic acids, and fatty acids. Additionally, radish pink and radish white responded nearly similar with higher levels of all amino acids except aspartic and glutamic acids. The amines, hydroxylamine, oxoproline, and methyl alaninate were detected in relatively high amounts in mustard, pak choi, and radish pink, whereas radish white had low levels in their microgreens.

Lastly, metabolite profiling also detected significant levels of polyphenol sinapinic acid in the microgreens, with elevated levels in mustard and pak choi, followed by radish white and radish pink.

### 3.6. The fatty acid profile of *Brassicaceae* microgreens showed beneficial lipids

The FAMEs (fatty acid methyl esters) based profiling in *Brassicaceae* microgreens identified six primary fatty acids (saturated and unsaturated). Saturated fatty acids include palmitic acid, and stearic acid, whereas unsaturated fatty acids include oleic acid, linoleic acid, eicosenoic acid, and erucic acid (Figure 4). Additionally, two terpenes, neophytadiene, and phytol, were also observed. The relative abundances of palmitic acid, stearic acid, oleic acid, and linoleic acid were similar among the microgreens. However, differences in the levels of erucic acid and phytol were observed. Erucic acid was less in radish pink and highest in radish white.

**Figure 4.**
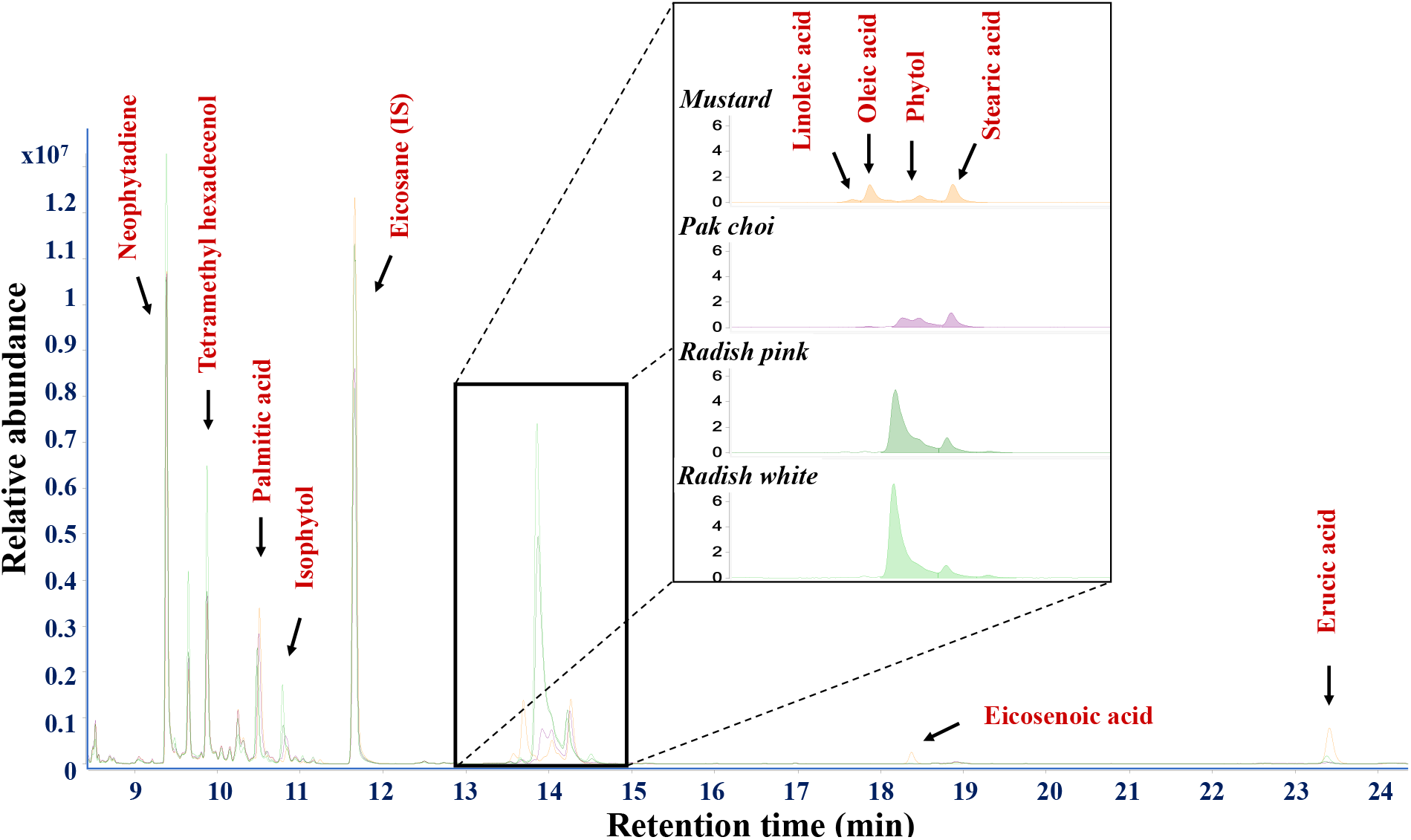
*Brassicaceae* microgreens have a beneficial lipid composition in their fatty acid profile. The profiling using FAMEs (fatty acid methyl esters) revealed the presence of saturated and unsaturated fatty acids. The orange, purple, dark green, and light green GC-FAMES spectra of mustard, pak choi, radish pink, and radish white, respectively. Eicosane (IS) was used as an internal standard at a concentration of 0.01 %.

The presence of essential fatty acids and terpenes identified through GC-MS FAMEs, i.e., oleic acid, linoleic acid, neophytadiene, and phytol makes *Brassicaceae* microgreens nutritionally rich (Figure 4,5). These unsaturated fatty acids are not synthesized in the human body and should be involved in diet. They play several roles in human nutrition, including the building block of the cell, cell membrane, hormone production, blood pressure regulation, inflammatory responses, etc. (Chen and Liu, 2020). Besides the essential fatty acids, average erucic acid levels were less than 5 % in all the *Brassicaceae* microgreens samples, which are considered safe for human ingestion (Figure 5). Overall, the fatty acid profiles in microgreens are encouraging, given that many are nutritionally essential.

**Figure 5.**
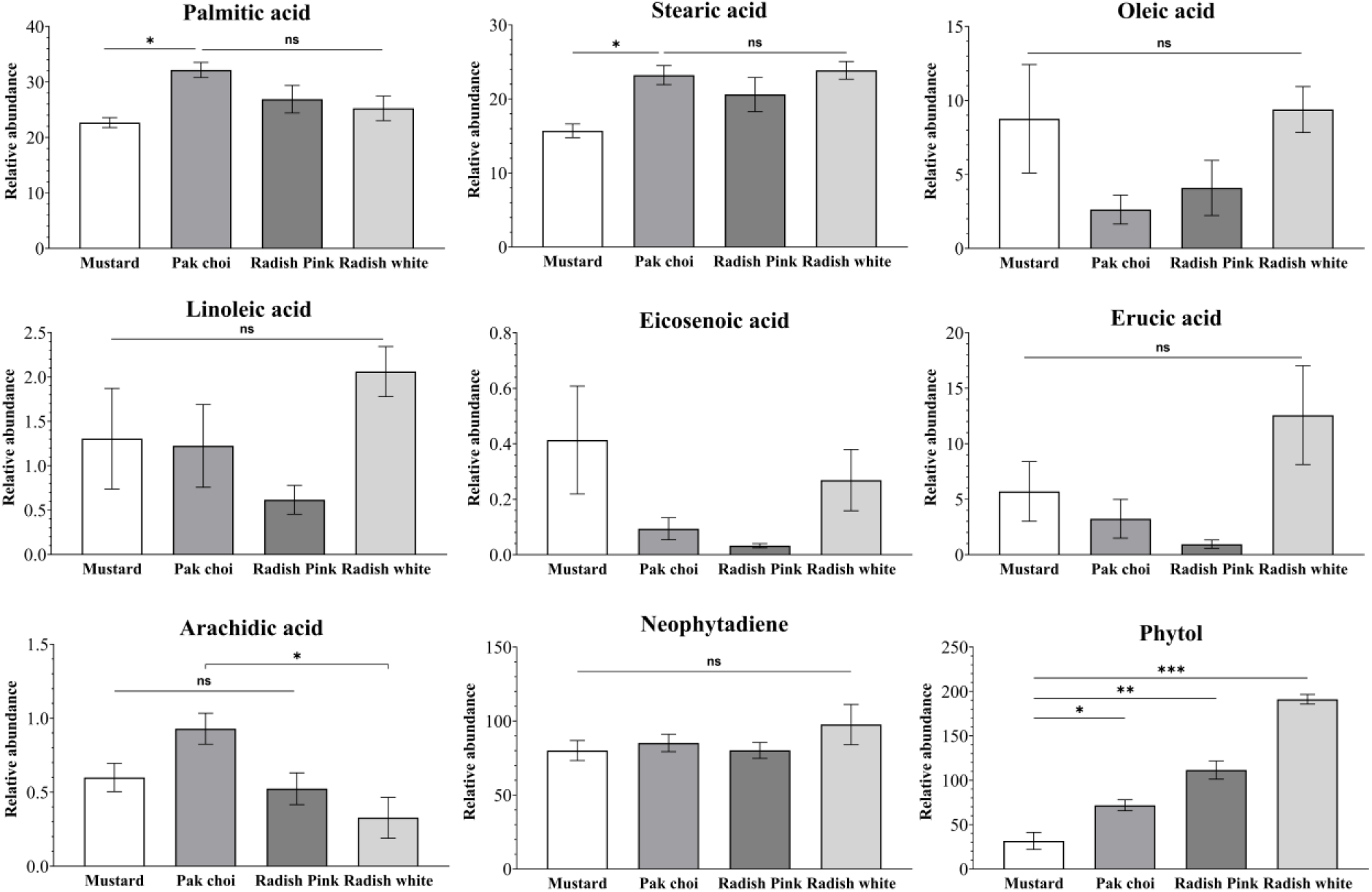
Qualitative levels of primary fatty acids and terpenes in *Brassicaceae* microgreens identified through GC-MS. The relative peak area abundance was normalized with the internal standard Eicosane and expressed as mean ± standard deviation (n=3). The abbreviation ‘ns’ denotes the absence of a statistically significant difference between the replicates.

## 4. Conclusion

Microgreens are the edible young seedlings of various vegetables, herbs, and flowers consumed at the two-leaf stage. These are reported to be rich in essential nutrients and other health beneficial components for being considered as functional food. The current study evaluates their dietary potential through biochemical, minerals, metabolite, and fatty acid profiles. The *in vitro* cultivation under controlled growth conditions was best achieved for microgreens of mustard (*Brassica juncea*), pak choi (*Brassica rapa subsp. chinensis*), radish pink (*Raphanus sativus*), and radish white (*Raphanus ruphanistrum*). The suitable substrate composition for optimum growth of *Brassicaceae* microgreens was optimized, and the 1:1 coco-peat and vermiculite ratio was observed to be the best for their germination and growth. The biochemical and mineral composition analysis (using ICP-MS) confirmed their promising nutritional, antioxidant nature and as excellent sources of minerals. While the carbohydrate, lipids levels in microgreens were comparable to mature *Brassicaceae* leaves, the total proteins were lower. The total polyphenols in microgreens were higher than in mature plants, indicating that including them in the diet can provide healthy human nutrition. Mineral profiling exhibited promising levels of Fe, Mn, Mg, K, and Ca in microgreens that are of health relevance. Further, when the mineral levels in *Brassicaceae* microgreens were compared to recommended dietary allowance, it is clear that they can meet a significant proportion of daily nutrient needs and assist in overcoming “hidden hunger.” The GC-MS based metabolite profiles of microgreens identified several small molecules belonging to sugars, amino acids, fatty acids, organic acids, polyphenols, sugar alcohols, and amines with nutritional significance. The FAMEs-based profiles show the presence of essential fatty acids, oleic acid, linoleic acid, and terpenes that are considered nutritionally important. All these nutritional parameters show that the *Brassicaceae* microgreens have health-beneficial role and can be used as excellent nutritive sources as functional food. Recent studies have shown that UV-B irradiation in plants can enhance their polyphenol levels. In future studies, such conditions can be optimized in microgreens to develop biofortified food.

## ACKNOWLEDGMENT

SKM acknowledges Science and Engineering Research Board (SERB) Early career research funding (File No: ECR/2016/001176). YP acknowledges the SERB, Ministry of Education, and IIT Mandi for PhD fellowship.

## Declarations

All the authors declare no conflict of interest.

## Notes

### Competing Interest Statement

The authors have declared no competing interest.

