## Supplementary file for "Metabolic, Biochemical, Mineral and Fatty acid profiles of edible *Brassicaceae* microgreens establish them as promising functional food"

| | | |
|---|---|---|
| **Table S1.** | Metabolites detected in the aqueous methanol soluble extracts of *Brassicaceae* microgreens using GC-MS | **Page 2** |
| **Table S2.** | Absolute abundances of Macrominerals and microminerals in various *Brassicaceae* microgreens | **Page 3-4** |
| **Figure S1.** | The relative abundances of (A) sugars and (B) amino acids in *Brassicaceae* microgreens | **Page 5** |
| **Figure S2.** | The relative abundances of metabolites among the *Brassicaceae* microgreens | **Page 6-7** |

.

**Table S1. Metabolites detected in the aqueous methanol soluble extracts of *Brassicaceae* microgreens using GC-MS**. The extracts are derivatised with MSTFA. The identities of the metabolite peaks are based on the fragments (m/z) and similarity scores against available in-house standards, NIST and Fiehn library.

| S.No. | RT (min) | Metabolite Name | Compound class | m/z | Score (%) |
|---|---|---|---|---|---|
| 1 | 12.06 | Lactic Acid, 2TMS derivative | Organic acids | 147, 73, 117, 148, 190 | 96 |
| 2 | 12.28 | Glycolic acid, 2TMS derivative | Organic acids | 147, 73, 177, 148, 66 | 93 |
| 3 | 12.75 | L-alanine | Amino acids | 147, 116, 73, 204, 148 | 91 |
| 4 | 12.91 | Hydroxylamine, 3TMS derivative | Other bioactive compounds | 73, 133, 146, 119, 147 | 97 |
| 5 | 14.52 | L-Valine, 2TMS derivative | Amino acids | 144, 73, 147, 218, 145 | 97 |
| 6 | 14.93 | Benzoic Acid, TMS derivative | Organic acids | 179, 135, 105, 77, 180 | 97 |
| 7 | 15.27 | Ethanolamine, 3TMS derivative | Other bioactive compounds | 174, 73, 147, 100, 86 | 98 |
| 8 | 15.34 | L-Leucine, 2TMS derivative | Amino acids | 158, 73, 159, 147, 102 | 95 |
| 9 | 15.39 | Glycerol, 3TMS derivative | Other bioactive compounds | 73, 147, 205, 133, 117 | 86 |
| 10 | 15.45 | 4-Dimethylamino-2(5H)-furanone | Other bioactive compounds | 127, 69, 99, 71, 56 | 74 |
| 11 | 15.66 | DL-isoleucine | Amino acids | 158, 73, 218, 147, 159 | 96 |
| 12 | 15.78 | 2-Butenedioic acid, (Z)-, 2TMS derivative | Organic acids | 147, 73, 75, 148, 45 | 87 |
| 13 | 15.86 | Glycine, 3TMS derivative | Amino acids | 174, 73, 147, 175, 248 | 97 |
| 14 | 16.21 | Glyceric acid, 3TMS derivative | Organic acids | 73, 147, 189, 292, 133 | 96 |
| 15 | 16.48 | Nonanoic acid, TMS derivative | Organic acids | 215, 117, 73, 75, 129 | 87 |
| 16 | 16.61 | Serine, 3TMS derivative | Amino acids | 204, 73, 218, 147, 100 | 97 |
| 17 | 16.99 | L-Threonine, 3TMS derivative | Amino acids | 218, 219, 291, 220, 293 | 89 |
| 18 | 17.26 | L-Aspartic acid | Amino acids | 174, 73, 100, 70, 75 | 92 |
| 19 | 18.15 | Timonacic, 2TMS derivative | Other bioactive compounds | 73, 160, 161, 74, 162 | 82 |
| 20 | 18.29 | Malic acid, 3TMS derivative | Organic acids | 233, 147, 73, 245, 189 | 95 |
| 21 | 18.40 | 3-Amino-2-piperidone, 2TMS derivative | Other bioactive compounds | 100, 115, 243, 128, 73 | 79 |
| 22 | 18.55 | Methyl L-alaninate, 2TMS derivative | Amino acids | 188, 84, 73, 189, 156 | 70 |
| 23 | 18.59 | Erythritol, 4TMS derivative | Organic acids | 217, 103, 205, 117, 133 | 91 |
| 24 | 18.71 | L-5-Oxoproline, , 2TMS derivative | Other bioactive compounds | 156, 73, 147, 157, 258 | 90 |
| 25 | 18.79 | 4-Aminobutanoic acid, 3TMS derivative | Organic acids | 174, 73, 147, 175, 75 | 91 |
| 26 | 19.85 | L-Glutamic acid, 3TMS derivative | Amino acids | 246, 73, 128, 147, 247 | 96 |
| 27 | 19.98 | Phenylalanine, 2TMS derivative | Amino acids | 73, 218, 147, 100, 219 | 95 |
| 28 | 20.49 | L-asparagine | Amino acids | 73, 116, 75, 141, 132 | 95 |

| | | | | | |
|---|---|---|---|---|---|
| 29 | 20.52 | D-Arabinose | Sugars | 103, 217, 307, 129, 189 | 87 |
| 30 | 20.68 | D-(+)-Ribono-1,4-lactone (R,R,R)-, 3TMS derivative | Other bioactive compounds | 73, 147, 117, 75, 133 | 79 |
| 31 | 21.36 | Ribitol, 4TMS derivative (Internal standard)) | Sugars | 217, 205, 218, 189, 204 | 95 |
| 32 | 21.42 | meso-Erythritol, 4TMS derivative | Sugars | 217, 205, 218, 189, 204 | 86 |
| 33 | 21.88 | L-glutamine | Amino acids | 73, 156, 147, 155, 75 | 91 |
| 34 | 22.45 | L-Threitol, 4TMS derivative | Sugars | 147, 103, 117, 189, 133 | 74 |
| 35 | 22.52 | Metabolite@22.52 | - | 221, 429, 355, 207, 281 | 84 |
| 36 | 22.77 | Citric acid, 4TMS derivative | Organic acids | 73, 147, 273, 75, 274 | 96 |
| 37 | 22.90 | Myristic acid, TMS derivative | Fatty acids | 117, 129, 285, 132, 43 | 79 |
| 38 | 24.01 | D-Fructose | Sugars | 73, 103, 147, 307, 218 | 96 |
| 39 | 24.32 | d-Mannose | Sugars | 73, 147, 205, 309, 160 | 82 |
| 40 | 24.72 | L-Lysine, 4TMS derivative | Amino acids | 73, 156, 174, 128, 317 | 87 |
| 41 | 25.04 | d-Glucose | Sugars | 73, 147, 205, 103, 160 | 97 |
| 42 | 25.14 | L-Tyrosine, 3TMS derivative | Amino acids | 218, 219, 100, 280, 220 | 94 |
| 43 | 25.44 | Ethyl α-D-glucopyranoside, 4TMS derivative | Sugars | 204, 73, 147, 205, 217 | 91 |
| 44 | 27.68 | Palmitic Acid, TMS derivative | Fatty acids | 117, 73, 313, 75, 129 | 96 |
| 45 | 30.64 | Myo-Inositol, 6TMS derivative | Other bioactive compounds | 73, 217, 305, 129, 204 | 93 |
| 46 | 33.75 | α-Linolenic acid, TMS derivative | Fatty acids | 73, 75, 79, 67, 95 | 85 |
| 47 | 34.49 | Stearic acid, TMS derivative | Fatty acids | 117, 75, 129, 341, 43 | 95 |
| 48 | 34.73 | Sinapinic acid, 2TMS derivative | Poly-phenol | 75, 338, 368, 323, 339 | 90 |

**Table S2. Absolute abundances of Macrominerals and microminerals in various *Brassicaceae* microgreens.** The abundances quantified in this current study and other studies from literature are tabulated. (NA- Data not available)

| Macro-minerals (mg/g DW) | | | | | | | | |
|---|---|---|---|---|---|---|---|---|
| | **Kyriacou *et al* 2021** (Pak choi) | **Mezeyova *et al* 2022** (Radish) | **Weber 2017** (Broccoli) | **Samuoliene *et al* 2019** (Broccoli) | **This study** | | | |
| | | | | | Mustard | Pak choi | Radish pink | Radish white |
| **Na** | 2.91 | 3.29 | 65 | NA | 20.7 | 26.3 | 20.9 | 18.16 |
| **Mg** | 4.8 | 8.94 | 4 | 4.54 | 7.35 | 8.59 | 6.5 | 6.85 |
| **K** | 17.05 | NA | 9 | NA | 56.28 | 58.29 | 52.94 | 38.66 |
| **Ca** | 9.93 | 4.06 | 4 | 14.3 | 14.39 | 16.91 | 10.37 | 9.35 |
| **Trace-minerals (μg/g DW)** | | | | | | | | |
| **Mn** | 149 | 32.3 | 30 | NA | 115.7 | 88.5 | 51.2 | 50.4 |
| **Fe** | 190 | 105.8 | 70 | 350 | 1021 | 266.3 | 624.6 | 293.6 |
| **Cu** | NA | NA | 10 | NA | 7.43 | 9.6 | 8.02 | 6.7 |
| **Zn** | 143 | 43.7 | 70 | NA | 135.89 | 19.6 | 13.6 | 11.9 |

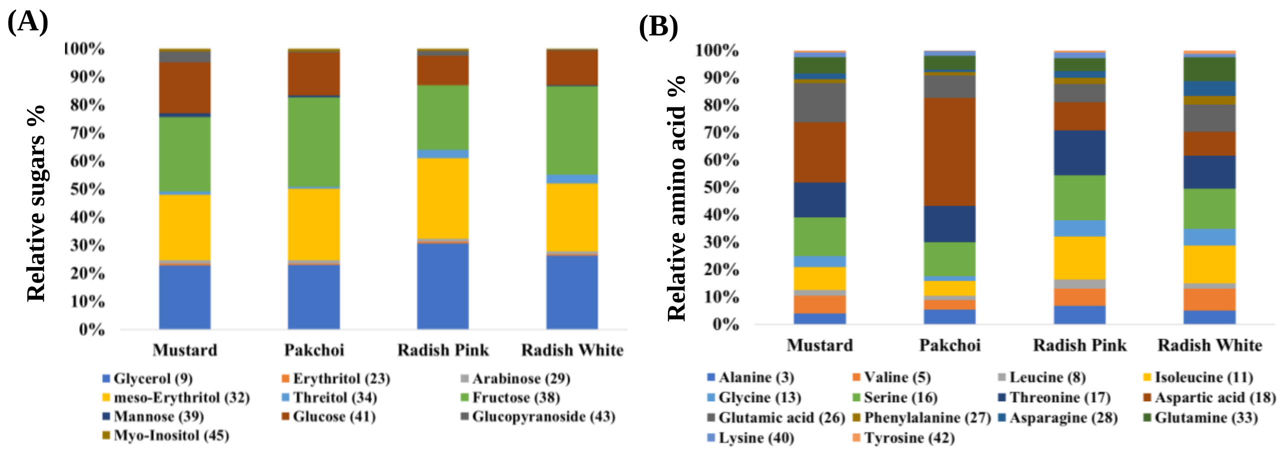


**Figure S1. The relative abundances of (A) sugars and (B) amino acids in *Brassicaceae* microgreens.** The stacked plots are based on metabolite peak areas analysed from GC-MS (from Table S1).

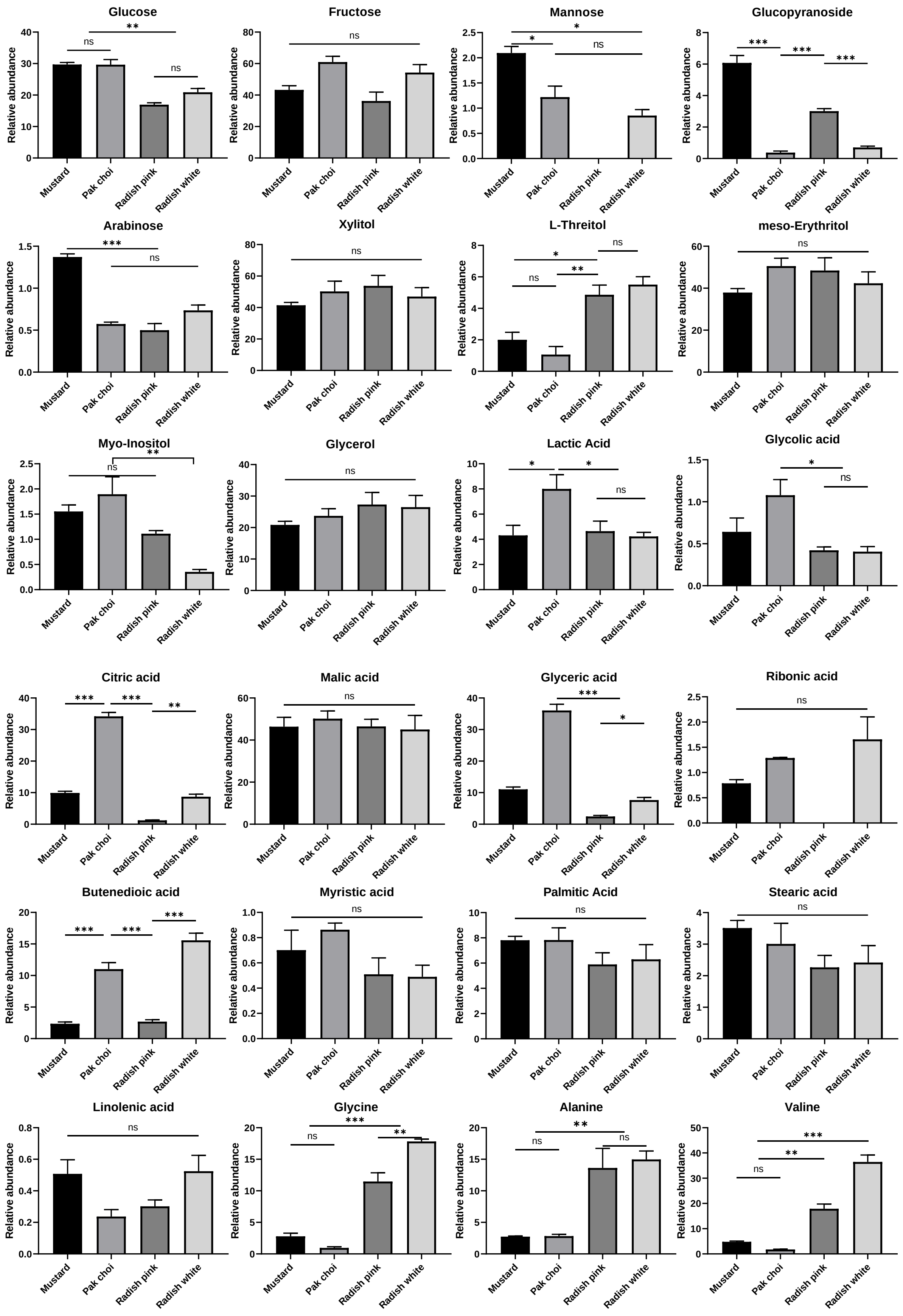
Glucose
Fructose
Mannose
Glucopyranoside
Arabinose
Xylitol
L-Threitol
meso-Erythritol
Myo-Inositol
Glycerol
Lactic Acid
Glycolic acid
Citric acid
Malic acid
Glyceric acid
Ribonic acid
Butenedioic acid
Myristic acid
Palmitic Acid
Stearic acid
Linolenic acid
Glycine
Alanine
Valine
Relative abundance
Mustard
Pak choi
Radish pink
Radish white
ns
*
**
***

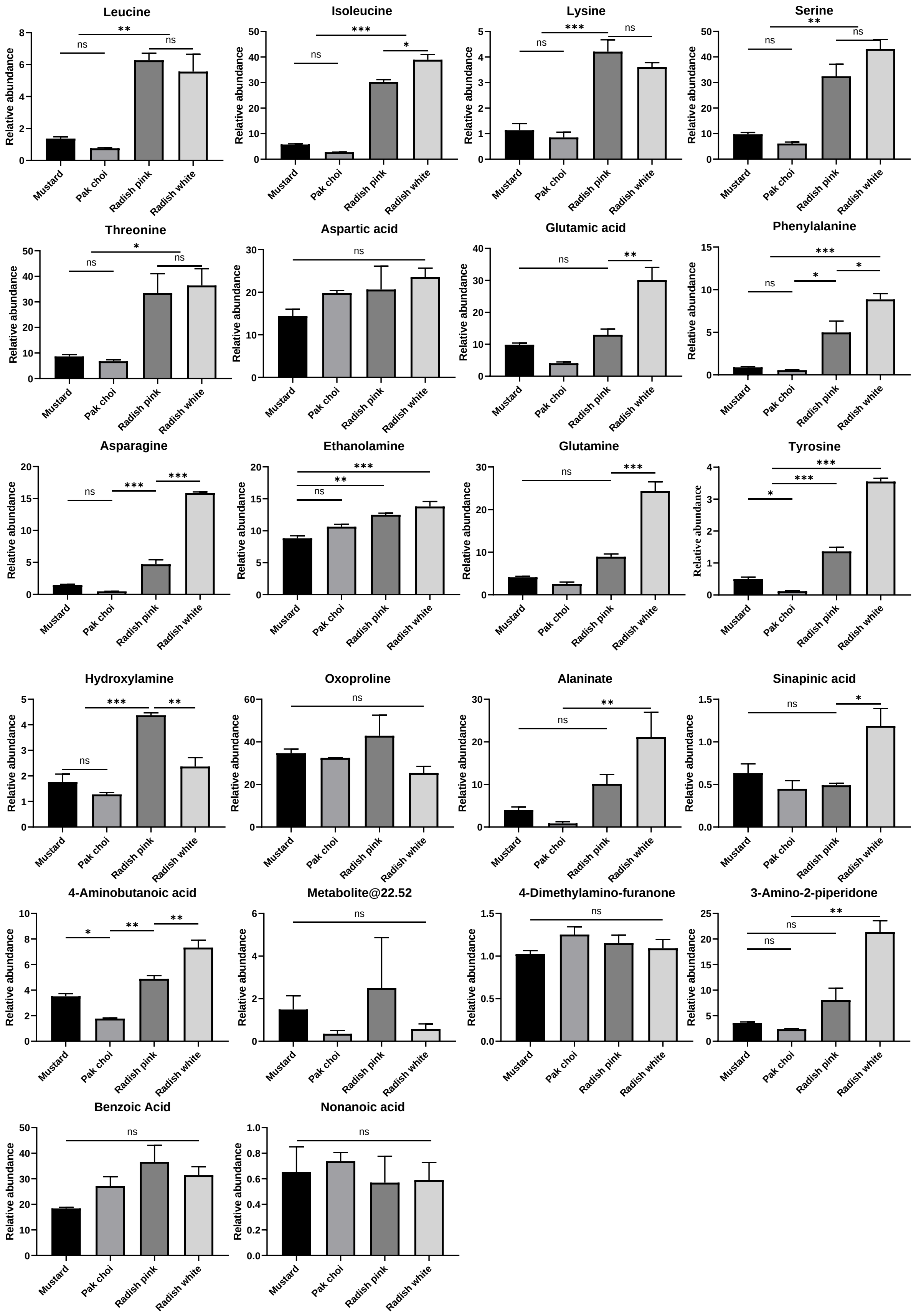


**Figure S2.** The relative abundances of metabolites among the *Brassicaceae* microgreens analysed through GC-MS.
